# varCADD: large sets of standing genetic variation enable genome-wide pathogenicity prediction

**DOI:** 10.1101/2024.09.24.614666

**Authors:** Lusiné Nazaretyan, Philipp Rentzsch, Martin Kircher

## Abstract

Machine learning and artificial intelligence are increasingly being applied to identify phenotypically causal genetic variation. These data-driven methods require comprehensive training sets to deliver reliable results. However, large unbiased datasets for variant prioritization and effect predictions are rare as most of the available databases do not represent a broad ensemble of variant effects and are often biased towards protein-coding genome, or even towards few well-studied genes. To overcome these issues, we propose several alternative training sets derived from subsets of human standing variation. Specifically, we use variants identified from whole-genome sequences of 71,156 individuals contained in gnomAD v3.0 and approximate the benign set with frequent and the deleterious set with rare standing variation. We apply the Combined Annotation Dependent Depletion framework (CADD) and train several alternative models using CADD v1.6. Using the NCBI ClinVar validation set, we demonstrate that the alternative models have state-of-the art accuracy, outperforming the widely used pathogenicity score CADD v1.6 in certain genomic regions.

Being larger than conventional databases, including the evolutionary-derived training dataset of about 30 million variants in CADD, standing variation datasets cover a broader range of genomic regions and rare instances of the applied annotations. For example, they cover more recent evolutionary changes common in gene regulatory regions, which are more challenging to assess with conventional tools. Finally, datasets derived from standing variation better represent allelic changes in the human genome and do not require extensive simulations and adaptations to annotations of the evolutionary-derived sequence alterations used for CADD training. We provide datasets as well as trained models to the community for further development and application.

**Suggestion for a Graphical Abstract:** 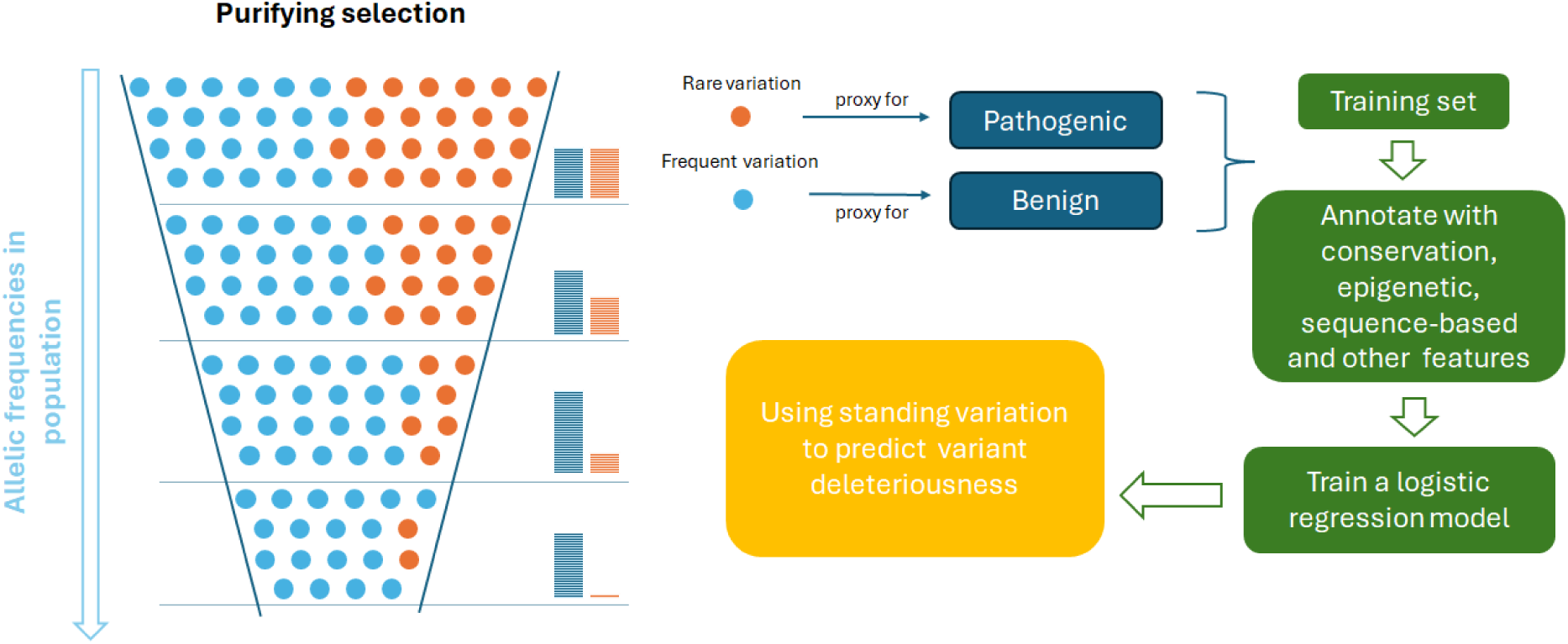

**Author‘s Summary:** Here, we are presenting the varCADD approach for predicting variant deleteriousness. Throughout time, pathogenic allelic changes are selected against by purifying selection, while neutral or beneficial changes can be passed along to next generations. Consequently, the frequencies of pathogenic variants are decreasing, beneficial alleles are increasing and frequencies of neutral variants are subject to drift. For that, allele frequencies in standing variation can be used as a proxy for their deleteriousness. To train a machine learning model for variant prioritization, frequent variants from gnomAD 3.0 were used as proxy-benign set and rare variants as proxy-deleterious set. The resulting training set exceeds excisting data sets in their size and allows for genome-wide coverage of molecular effects. The training set was annotated with sequence conservation, epigenetic, sequence-based and other features using the CADD v1.6 framework, after which a logistic regression model was trained. The output of the model can be interpreted as a probability for a variant to have a deleterious effect on genetic function.

## Introduction

In recent years, substantial advancements have been made in identifying phenotypically causal variants across the human genome. Machine learning-based models for prioritizing these variants have played a crucial role in this process. However, the application of machine learning on genomic data often faces challenges as comprehensive and unbiased training data is rare (Shihab et al. 2015; Quang, Chen, and Xie 2015). Many collections of pathogenic and benign variants are biased towards the protein-coding genome, specifically the identification of missense (i.e., amino acid substitution) variants (Ellingford et al. 2022; Spielmann and Kircher 2022). There is no broad ascertainment of clinical variants, and few well-studied genes tend to be overrepresented in these sets (Sundaram et al. 2018). Further, most of the annotated pathogenic variants have strong detrimental effects, missing out variants with low but significant functional impact or those with complex pathways of functional impact (Ciesielski et al. 2024; Castel et al. 2018; Domingo, Baeza-Centurion, and Lehner 2019; Virolainen et al. 2022; Hibbs et al. 2009). Besides, datasets often contain only a limited number of variants of different molecular effects or might have mixed quality of variant interpretation (Harrison et al. 2017; Lek et al. 2016). It is therefore challenging to train machine learning models of variant pathogenicity, especially genome-wide, across different molecular processes and effect sizes.

To overcome some of these issues, the Combined Annotation Dependent Depletion (CADD) framework (Kircher et al. 2014) introduced an alternative set of proxy-deleterious and proxy-neutral variants to train machine learning models for ranking all kinds of potentially disease causal variants from genome sequencing (i.e., single nucleotide and insertion/deletion variants, as well as coding and non-coding regions of the genome). Rather than directly predicting variant pathogenicity, CADD approximates pathogenicity by modeling deleteriousness, i.e. variants that might be excluded by purifying selection during species evolution. The proxy-neutral set consists of around 14 million human-lineage-derived sequence alterations which saw many generations of purifying selection and is therefore assumed to be a proxy for neutral variation. The proxy-deleterious variation is a set of simulated variants obtained by a custom genome-wide simulator of germline variation. It simulates ‘de novo’ variants according to the substitution frequencies and insertion/deletion (InDel) lengths observed in the proxy-neutral set, accommodating a local (megabase resolution) adjustment of mutation rates and asymmetric CpG-specific mutation rates. Using this data, CADD trains an unbiased learner that generalizes well to the variation in the entire human genome, outperforms other genome-wide approaches and still performs on-par or even better than highly specialized pathogenicity scores (Wu et al. 2021; Rentzsch et al. 2019).

Since the first publication of CADD, negative correlation has been observed between its scores and the frequency of variants from the 1000 Genomes Project (Kircher et al. 2014). This observation is in line with studies demonstrating an inverse relationship between the effect size of a variant and its frequency in the population (Gudmundsson et al. 2022; Cirulli and Goldstein 2010; Bomba, Walter, and Soranzo 2017; Park et al. 2011). Thus, the frequency of alleles with detrimental effects is controlled by purifying selection, which is the reason why this inverse relationship is skewed towards rare variants for traits strongly influenced by natural selection (Marth et al. 2011). In contrast, variants responsible for less constrained quantitative phenotypes such as height or eye color, as well as variants underlying late-onset diseases, may be more common due to weaker selection effects (Lettre 2014). This suggests that by contrasting alleles of different frequencies one can also gain insights into functional effects of variation in the human genome.

Leveraging allele frequency to gauge the functional impact of genetic variation necessitates extensive datasets of genome-wide variant data to comprehensively cover molecular effects and accurately estimate variant frequencies. When CADD was developed in 2014, variant simulations and human-species derived sequence alterations were used for its development due to the lack of large-scale genome sequencing data and variant sets. However, reduced sequencing costs have enabled comprehensive genomic datasets from thousands of individuals, facilitating the examination of the effects of variation within both coding and noncoding regions of the genome (Gudmundsson et al. 2022). Initiatives like gnomAD (S. Chen et al. 2022), TOPmed (Taliun et al. 2021), and ALFA (Phan et al. 2020) provide openly accessible variant data sets from thousands of individuals. While the dataset from the 1000 Genomes Project contained around 88 million variants (The 1000 Genomes Project Consortium et al. 2015), gnomAD 3.0 includes around 602 million single nucleotide variants (SNVs) and 105 million InDel variants (S. Chen et al. 2022). The growing number of sequenced genomes from various initiatives is poised to enhance the interpretation of genetic variants. Larger and more broadly sampled genomic dataset not only encompasses a greater abundance of rare variants but also refine our ability to accurately estimate allele frequencies between populations or geographic ancestries.

In this manuscript, we investigate the potential of using variant data for the purpose of training genome-wide models for variant prioritization. We construct several alternative training sets based on standing variation, using frequent variants as a proxy for (more) neutral and rare variants as a proxy for (more) deleterious variation. We then employ logistic regression on each of these training set, the results of which demonstrate comparable performance with CADD. Moreover, augmenting the CADD training set of human-derived data with standing variation outperforms the original model in identifying specific types of pathogenic variants, such as stop-loss (i.e., non-sense) or inframe variation in coding regions, regulatory variants, 3’ and 5’ UTR variants, as well as other types of noncoding variation.

## Results

### Evidence of a proxy-deleterious state of low allele frequency variants

Initially, we assessed the relationship between CADD v1.6 deleteriousness scores (Rentzsch et al. 2019) and the allele frequency for the Genome Aggregation Database (gnomAD) v3.0 (S. Chen et al. 2022). This global cohort contains variants identified from genome sequencing of 71,156 individuals, excluding first or second-degree relatives, mapped to the GRCh38 genome build. We randomly sample gnomAD variants with PHRED-scaled CADD scores from 1 to 50, filtering out variants with zero allele count or allele frequency (AC=0 or AF=0). **Figure 1** shows the mean allele frequency for each bin of PHRED scores as well as the proportion of singletons in each bin. We observe an inverse relationship between the allele frequency and PHRED scores, i.e., the predicted deleteriousness of a variant in the gnomAD set is negatively correlated with its frequency. This relation is especially clear for the lower-scoring and largest part of the genome, while it is less stable for higher deleterious scores (i.e., the PHRED scores above 25 representing less than 0.3% of the genome). Considering variants across all PHRED bins, scaled CADD scores and allele frequencies show a Spearman correlation with allele frequencies of −0.116 (n = 3,264,650, p-value < 1.171 * 10^−157^) or −0.131 (n = 1,443,979, p-value < 1.13*10^−187^) when excluding singletons.

**Figure 1.**
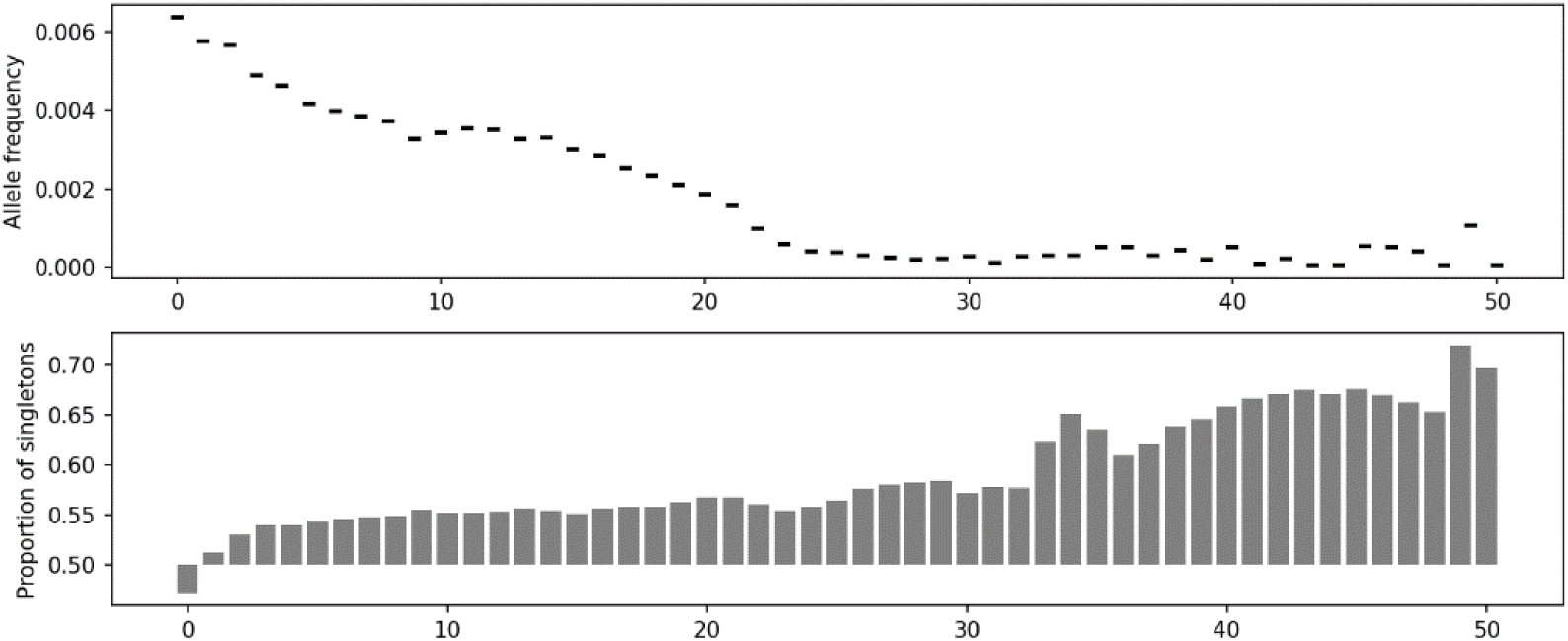
Inverse correlation between deleteriousness scores and allele frequencies. Average allele frequency for bins of PHRED-scaled CADD scores up to 50 for variants sampled from the gnomAD v3.0 database. For each bin of PHRED scores up to 100,000 variants were sampled from gnomAD v3.0. Starting from PHRED score 29, the number of available variants with the corresponding score starts to decrease sharply. For example, for 49 < PHRED ≤ 50, only 581 variants were available. At the same time, the number of singletons is increasing with PHRED score, which makes conclusions based on the right tale of the distribution less certain. In general, average allele frequency decreases with PHRED scores indicating that there is an inverse relationship between predictions of deleteriousness derived from CADD and average allele frequency.

Studies using genome and exome sequencing data allow quantitative assessment of purifying selection. For example, Dukler et al. (2022) estimate that ∼0.4–0.7% of the human genome is ultra-selected, which suggests 0.26–0.51 strongly deleterious *de novo* mutations per generation. Moreover, singletons, variants observed only once in the entire cohort and potentially harboring *de novo* variation, are more likely to be located in functionally important loci (Ke, Taylor, and Cardon 2008). However, new mutations are challenging to assess as they are hard to detect and have fitness effects that are difficult to measure (Dukler et al. 2022). Nonetheless, various studies demonstrate that singletons, ultra-rare and rare variants accumulate in certain disease cohorts (Yang et al. 2022; Kryukov, Pennacchio, and Sunyaev 2007; Bomba, Walter, and Soranzo 2017; Momozawa and Mizukami 2021; Johansen et al. 2012; Griswold et al. 2015). This evidence along with the highly significant correlation of CADD scores with allele frequencies supports the idea that information about allele frequency can be leveraged further, delivering new biological insights and improving variant prioritization.

### Composition of alternative datasets

Capitalizing on the idea of purifying selection, we want to use variation of different allelic frequencies as datasets for training machine learning models for variant prioritization. For that, we continue with gnomAD v3.0. Due to the limited or questionable coverage of downstream annotations used for alternative haplotypes, unplaced contigs and the mitochondrial genome, we drop these variants from further analysis. As a result, we obtained 525 million SNVs and 68 million InDels. Much of the variation considered (51.1% of SNVs and 41.3% of InDels) has an allele count of 1 (AC = 1, singleton) in the database, i.e. it is observed only in a single individual, on one of their haplotypes. These singleton variants could potentially constitute the largest group of a proxy-pathogenic variant set for model training.

Further, we define all variants below a minor allele frequency (MAF) of 0.1% as rare variants and regard them as a second potential proxy-pathogenic set (MAF < 0.1% & AC > 1). Thus, as proxy-benign set, we define frequent variants, i.e. those greater than 0.1%. Although there is no universal definition of how often a variant should be observed in a population to be called frequent, numbers between 0.1% and 1% have been used before. They are considered to be conservative frequency cutoffs for recessive (1%) and dominant (0.1%) diseases, respectively (Bamshad et al. 2011). Further, population genetic studies demonstrate that severe disease-causing variants have much lower frequency than these thresholds (Whiffin et al. 2017). Thus, a cutoff of 0.1% for the proxy-benign set seems to deliver a reasonable separation from proxy-pathogenic variants for our purpose.

As the number of all variants differs significantly throughout the groups, with “singletons” being the most abundant and “frequent” variants the least abundant, we sample all the “singleton” and “rare” variants to match the number of frequent variants, so that each potential training set contains 26,255,876 SNVs and 8,864,604 InDels (**Table 1**). We use both “singleton” and “rare” variant sets as proxy for deleterious variation and “frequent” variants as proxy for neutral variation.

**Table 1.** Number of frequent, rare and singleton variants in the gnomAD database as well as the human-derived set used for training of CADD v1.6. While most variants in the processed gnomAD v3.0 dataset are singletons, variants with MAF >= 0.001 contribute 5.9%, in absolute numbers more than the number of human-derived variants used as a benign set for training the CADD v1.6 model.

| Frequency | Definition | SNVs | InDels |
| --- | --- | --- | --- |
| Singletons | AC = 1 | 268,404,480 | 27,934,609 |
| Rare | MAF < 0.001 & AC = 1 | 229,184,030 | 30,416,715 |
| Frequent | MAF ≥ 0.001 | 26,255,876 | 8,864,604 |
| Human-derived | human-lineage-derived sequence alterations | 14,011,297 | 1,688,847 |

### Features of alternative models

The CADD v1.6 training dataset is composed of 1028 features (Rentzsch et al. 2019). Among them are the reference and the alternative allele, as well as the inferred transition/transversion state. We have observed that the substitution rates of the four nucleotide bases adenine (A), guanine (G), cytosine (C) and thymine (T) differ across the three allele frequency groups defined above. For example, the proportion of C-to-T substitution in the set of “frequent” variants is 40.1%, whereas in the “singleton” set it is only 29.8% (**Error! Reference source not found.**). These shifts in representation of reference and alternative alleles as well as transition/transversion class in these alternative training sets can potentially cause label leakage and misguide model training. As a result, a model might for example assign a lower deleteriousness score to a variant with C to G substitution without any biological reasoning, but only because it is more frequent in the proxy-neutral training set than in proxy-deleterious training set. Therefore, we matched the substitution rates across the training sets to those of the “frequent” set.

Another set of features used to train CADD is derived from variant density. This set constitutes a total of 180 features, e.g., distance or number of rare, frequent or singleton SNV in a certain window in the BRAVO variant database (Taliun et al. 2021) as well as combinations of these features with selected annotations from the feature set (see Section *Model training* in *Methods*). BRAVO and gnomAD variant frequencies as well as presence/absence states will for obvious reasons be correlated, rendering these annotations potentially harmful for an unbiased model training (due to another potential source of label leakage). Hence, we train all models including and excluding annotations of variant density and compare their results. Otherwise, we use all features of CADD v1.6 to train a logistic regression model for each of the dataset pairs listed in Table 2. Additionally, we train a model on the original CADD dataset (human-derived versus simulation) excluding features derived from variant density annotations to assess the impact of these annotations on models not based on standing variation.

**Table 2.** Datasets used for training CADD v1.6 and our alternative models. Each pair of datasets is trained twice, including and excluding the annotations of variant density, to assess how those influence model predictions. Additionally, a model is trained based on the original CADD training set (human-derived vs. simulated) excluding variant density features.

| Identifier | Proxy-neutral | Proxy-deleterious | Variant density |
| --- | --- | --- | --- |
| fr | frequent | rare | excluded |
| fr' | frequent | rare | included |
| fs | frequent | singleton | excluded |
| fs' | frequent | singleton | included |
| rs | rare | singleton | excluded |
| rs' | rare | singleton | included |
| v1-6 | human-derived | simulated | excluded |
| v1-6' (CADD v1.6) | human-derived | simulated | included |

### Training optimized alternative models

In terms of machine learning, it is important to understand two specific characteristics of training CADD models. For one, CADD trains a surrogate model, i.e., its score assesses the putative deleteriousness of genomic variants, but it is applied to predict a wide range of variant effects, including pathogenicity, molecular function and variant impact. Furthermore, the training set of CADD has a substantial degree of mislabeling. For example, the set of simulated variants contains a high number of neutral variants, as simulated variants are “randomly” sampled across the genome, including regions of no functional constraint (Rentzsch et al. 2019). On the other hand, the proxy-benign set will contain functional changes that underly human-derived traits. For that, tuning and evaluation of CADD-like models is not sufficient on the holdout set of the training data, but needs external validation sets that cover various variant effects and are less prone to mislabeling.

Models for all current CADD releases (v1.4-1.7) have been trained using bulk logistic regression with 20 iterations. After each iteration, the model performance on external validation sets has been assessed and the 13^th^ model has been selected as the one with the best generalization (Schubach et al. 2024). To optimize alternative models, we follow the same scheme, training for 20 consecutive iterations. Due to the change in training data, we studied the effect on optimal training duration as well. For that, we assess and plot the models’ performance using Area Under the ROC (AUROC) curve and Area Under the Precision Recall Curve (AUPRC) after each iteration on a combination of 10 validation sets (see Section *Validation sets* in *Methods*). Based on the results, we identify the best iteration for each of the trained models (**Error! Reference source not found.**) and use them for all subsequent analyses. The selected iterations can be found in **Error! Reference source not found.**.

### State-of-the art models can be trained from standing variation

In the first step of the analysis, we want to assess whether the alternative models based solely on standing variation can achieve state-of-the-art performance for variant prioritization. For that, we compare the AUROC and AUPRC of newly trained models with that of CADD v1.6 on an independent validation set created from the NCBI ClinVar data (not used for model tuning). To create the dataset, we filtered SNVs and InDels with GRCh38 coordinates for their clinical significance benign, likely benign, pathogenic, and likely pathogenic. Further, we limited their length to a maximum of 50 base pairs. The resulting dataset contains 988,536 variants, 185,794 of which are pathogenic and 802,742 are benign.

**Figure 2** shows the ROC and PRC of trained models in comparison to CADD v1.6 on the NCBI ClinVar validation set. We observe that all models achieve a high classification rate, with the worst performing model in terms of recall having an AUROC of 0.953 (rs’, rare-singleton-with-variant-density) and the best performing models of 0.988 (fr’, frequent-rare-with-variant-density and fs’, frequent-singleton-with-variant-density). For precision scores, differences between models are rather small in most of the comparisons. The best alternative models are frequent-rare-with-variant-density (fr’) and frequent-singleton-with-variant-density (fs’), both slightly better than CADD v1.6 (recall 0.988 vs. 0984, precision 0.960 vs. 0.951). The worst models are rare-singleton with and without variant density annotations (rs’ and rs). We speculate that the composition of such training sets is not discriminating between variant effects enough, so that a model has more difficulties distinguishing between true pathogenic and benign variants properly. Across all models, performance differences were too small to derive a clear conclusion in favor or against one of the models. However, we note that alternative models based on frequent-rare and frequent-singleton sets show comparable performance to CADD v1.6.

**Figure 2.**
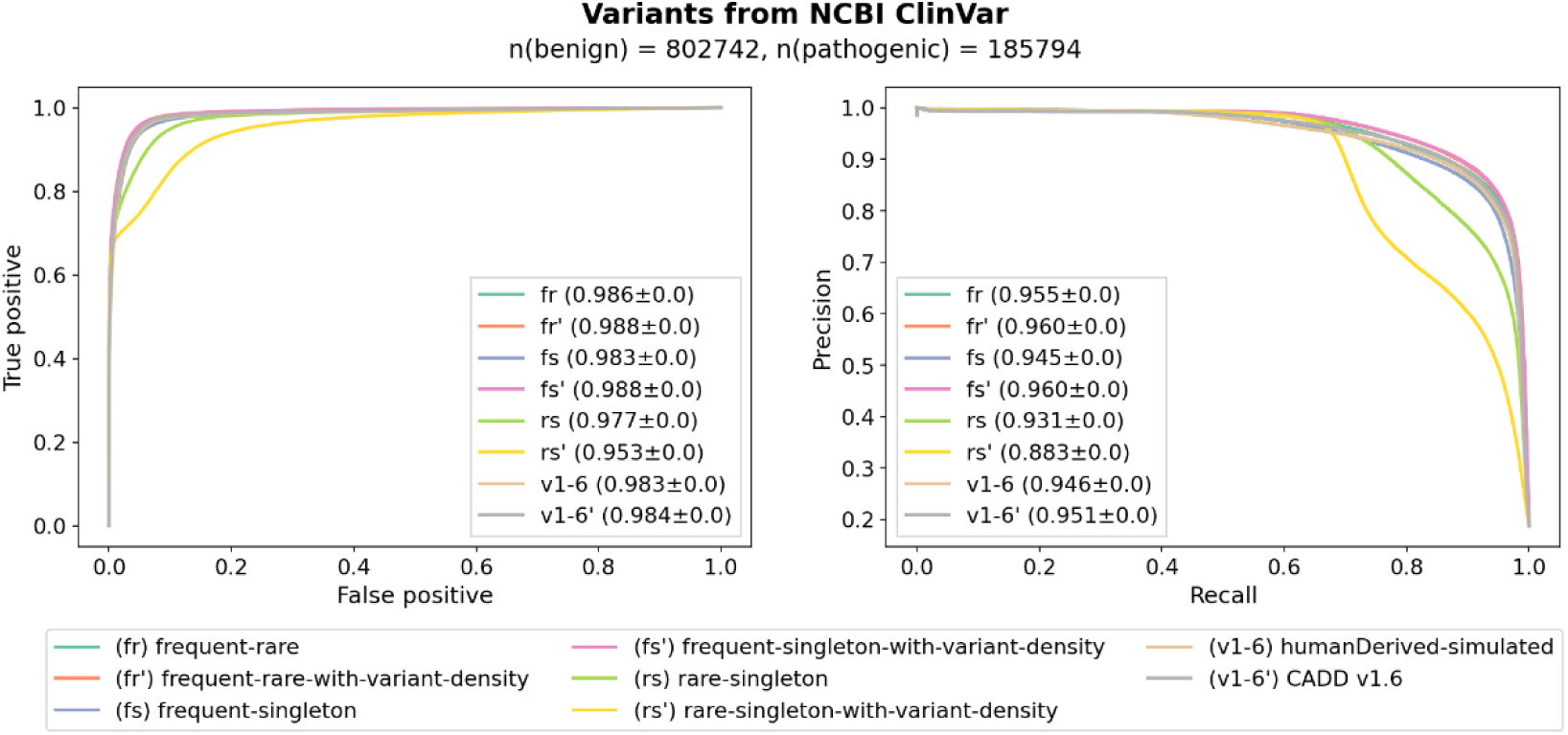
AUROC and AUPRC of trained models and CADD v1.6 on a validation set derived from NCBI ClinVar. The worst performing models are based on the rare vs. singleton datasets which might be explained by the fact that these proxies for pathogenic and benign sets are very close and thus more difficult to discriminate. The rest of the models have similar performance to CADD v1.6 and, hence, can be used to predict pathogenicity of genetic variation. Models that include variant density annotations are indicated with an apostrophe.

Further, the inclusion of variant density annotations seems to have a mixed effect: for models based on frequent-rare and frequent-singleton datasets, the performance seems to improve slightly; for models based on the rare-singleton set the opposite is the case. It is likely that annotations of variant density are beneficial for training sets with higher discriminative potential between pathogenic and benign sets because they can be used to distinguish these variants further, whereas in training data with higher label noise, these additional features and their noise hinders model training. Importantly, we do not observe a substantial label leakage due to these annotations as the performance of frequent-rare and frequent-singleton models on an external validation set does not deteriorate with their inclusion.

### Models have similar discriminative power for highly deleterious variants

While the overall performance seems similar across models, we wanted to understand whether there is a notable difference between model scores (**Figure 3**). We started our analysis by taking a closer look at a histogram/kernel density distribution of scores produced by different models calculated on predictions of 1 million potential SNVs randomly sampled from the human genome (**Figure 3A**). Here, two different distribution shapes can be distinguished. Models like rs (rare-singleton), rs’ (rare-singleton-with-variant-density) and fr (frequent-rare) have narrow shaped model score distributions (i.e. less variance) compared to all other models. This indicates that they do not exploit the entire score range, therefore, having lower resolution not only on the tails of the score distribution but also in the middle range. Further, the pathogenicity scores of these models are highly correlated (**Figure 3B**). Models trained with variant density features tend to have higher average scores and higher variance. This indicates that annotations of variant density have effects on model outcomes, shifting the average predicted pathogenicity to a higher (fr and fs) or lower value (rs and v1-6).

**Figure 3.**
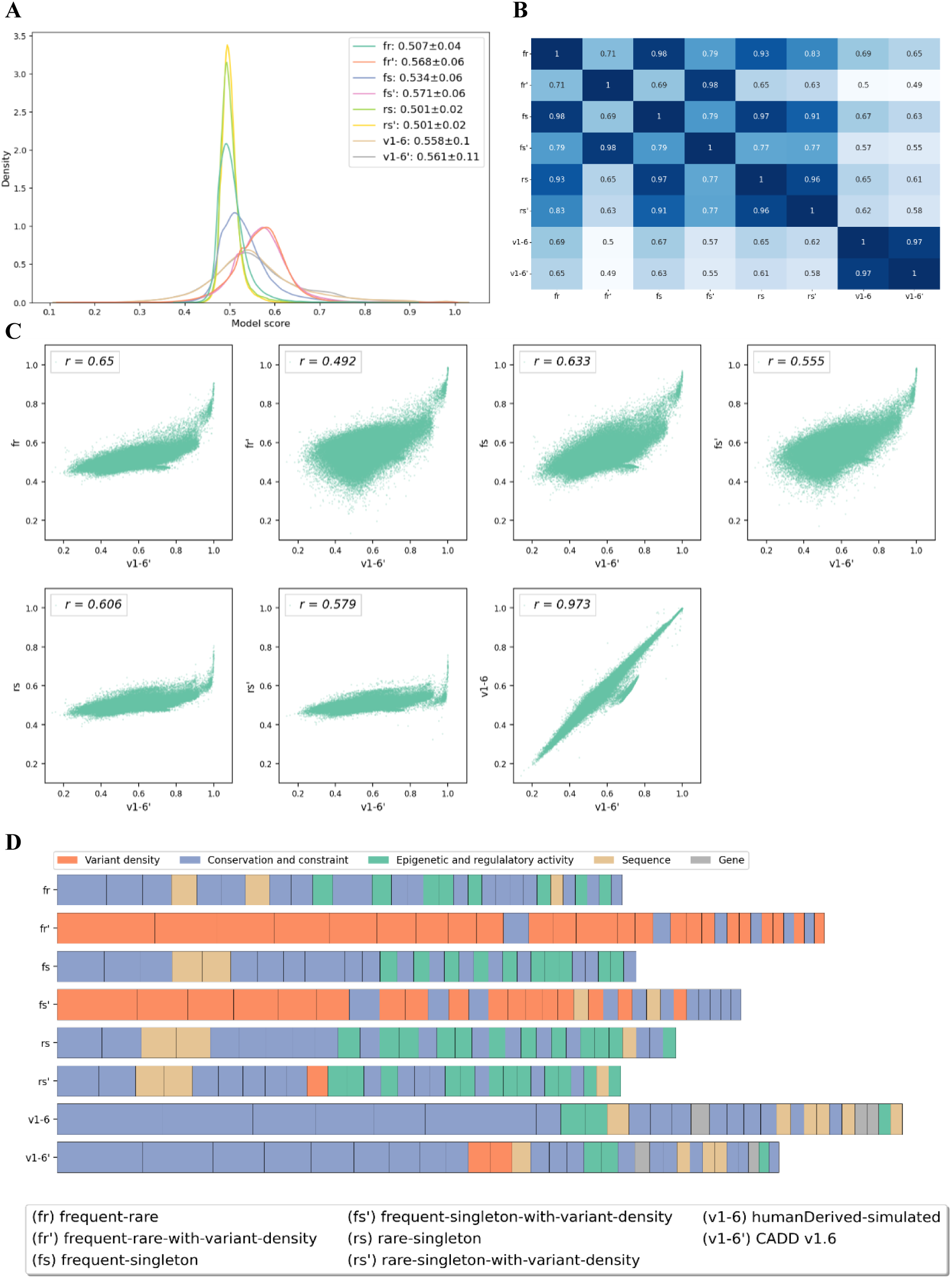
Distribution and correlation of scores as well as model coefficients of trained models. Panels A-C are based on a set of 1 million randomly sampled potential SNVs from the reference genome. Model labels are used as defined in Table 2. The label v1-6’ refers to the published CADD model, v1-6 is a model trained on the original CADD v1.6 data, but without the annotations of variant density. **A) Distribution of model scores.** Models based on sets of frequent-rare and rare-singleton variants do not exhibit major changes in the score distribution with inclusion of annotations of variant density. The model based on the set of frequent-singleton variants seems to be more sensitive to additional annotations. **B) Pearson correlation of model predictions.** Models fr’ (frequent-rare-with-variant-density) and fs’ (frequent-singleton-with-variant-density) are highly correlated with each other (Pearson rho = 0.98) and less correlated with their respective versions without annotations of variant density (fr:0.71, fs: 0.79). **C) Scatter plots of model predictions (y-axis) and CADD v1.6 scores (x-axis)**. Higher scores indicate higher predicted deleteriousness and higher probability of pathogenicity. Models tend to agree on highly pathogenic variants but diverge in scoring less deleterious variants. **D) Model coefficients of the first 30 most important features or feature combinations.** The features are color-grouped to facilitate the interpretation. The size of each block in a bar represents the absolute value of a model coefficient, the total length of a bar indicates the sum of coefficients. Conservation and constraint features are of high importance for almost all models, except for those based on frequent-rare and frequent-singleton sets trained with the full set of features (fr’ and fs’). For those, annotations of variant density dominate the model predictions.

Next, we analyzed the correlation of alternative models with CADD v1.6 scores. **Figure 3C** shows pairwise scatterplots of all models with CADD scores for the 1 million SNVs randomly selected above. A higher score indicates a higher predicted probability for a variant to be deleterious. While the CADD model trained without annotations of variant density (v1-6) is highly correlated (Pearson correlation coefficient of 0.97), the models trained on standing variation are only moderately correlated with CADD v1.6 scores (Pearson correlation coefficients from 0.49 to 0.65). The highest correlation is with the fr’ (frequent-rare-with-variant-density) and the lowest correlation is with fr. In the frequent-rare and frequent-singleton training sets, annotations of variant density seem to play an important role for score ranks. Importantly, most of the models tend to agree on the score of highly deleterious variants (according to CADD v1.6) but have different predictions when the functional impact of a variant might be more ambiguous. This is a known trend in scoring of variant deleteriousness as for example pathogenic variants have a significant impact that is easier to recognize. On the contrary, weaker or more quantitative effects are more difficult to capture and model, leading to disagreement between model predictions. This might also explain the only moderate correlations between models, as in the sampled set of 1 million variants the proportion of strongly deleterious variants is expected to be low, whereas variants with no or weak effects are expected to be most abundant.

### Alternative models do not overfit on variant density information

To further explore the differences in model predictions, we examine the coefficients of trained models as they can give an insight into the underlying causes of different score distributions and correlations. **Figure 3D** shows the assigned feature groups of the first 30 most important features in each model (**Error! Reference source not found.**). As feature values are normalized for all these models, coefficients can be compared directly.

In alternative models with variant density features, those are dominant for model decision. An exception here is the model rs’ (random-singleton-with-variant-density) where annotations of variant density have lower coefficients and are less frequent in the top 30 features. This observation supports the suspicion that singleton and rare variants are rather similar in their relation to variant density, having only minor differences in the annotations of variant density, so that model training cannot use them to separate the positive and the negative set. Contrary, for fr’ (frequent-rare-with-variant-density), 24 out of 30 features and, for fs’ (frequent-singleton-with-variant-density), 19 out of 30 features belong to the group of variant density, having also the highest coefficients. Hence, a bigger proportion of the variance explained by the model is attributed to the annotations of variant density.

Interestingly, in humanDerived-simulated, i.e., the original CADD models, conservation and constraint annotations are very dominant among the most important features (**Figure 3D**), whereas in models based on standing variation, annotations of epigenetic and regulatory activity as well as sequence-based features are of considerable importance. This might be explained by the consideration that standing variation contains more recent allelic changes with fewer generations of purifying selection as the evolutionary variants, and hence, are less-well separated by species conservation information. Consequently, the models trained on standing variation would put less weight on these features to distinguish between the proxy-pathogenic and proxy-benign variants in the training data.

### Substituting simulated variants for singleton variation in CADD

One of the drawbacks of CADD is its proxy-pathogenic training set based on a simplified simulation of *de novo* variants that might not reflect the entire complexity of natural selection and not perfectly reproduce the true distribution of variants across functional elements (Sabarinathan et al. 2016). For that, replacing the simulated data with real de novo variants might be advantageous for deleteriousness scoring. *De novo* variants are identified using trio-sequencing as each human carries up to 80 *de novo* variants (Sasani et al. 2019; Jónsson et al. 2017; Ségurel, Wyman, and Przeworski 2014). To replace the CADD dataset of 14 million simulated SNVs, one would need around 175,000 trios. However, trio studies are often elaborate and expensive, so that even with the largest initiatives of trio sequencing like the Simons Powering Autism Research (SPARK, 50,000 trios) (Wright et al. 2023; Feliciano et al. 2018) or the Deciphering Developmental Disorders (DDD, 14,000 trios) study (Wright et al. 2023; Feliciano et al. 2018), the amount of available data is not (yet) sufficient to replace the proxy-pathogenic set of CADD. Singletons can serve as an approximation for *de novo* variants as these rare variants also contain many newly occurring variants that have seen little purifying selection. For that, we replace the simulated data in the original CADD training set with the singleton set derived from the gnomAD v3.0 database and train another CADD-like model.

As the singleton set contains a much higher number of variants than the human-derived set (**Table 1**), we train balanced (hs for humanDerived-singleton) and unbalanced (hs-all for humanDerived-singleton-all) models (**Table 3**). For the balanced model, a subsample of the singleton set is used (14,011,297 SNV and 1,688,847 InDel), while all singleton variants (268,404,480 SNV and 27,934,609 InDel) are contrasted with the human-derived set for the unbalanced model, where we increase the class weights of the human-derived variants respectively. We speculate that the higher number of training variants could lead to a better coverage of rare annotations as well as broader genomic regions, which might be beneficial for the generalization power of the final model. The models are trained for up to 50 iterations, as we suspected that large datasets might require more intensive training to achieve the best results. The final number of iterations is chosen based on validation sets (see Methods section *Training optimized alternative models*).

**Table 3.**
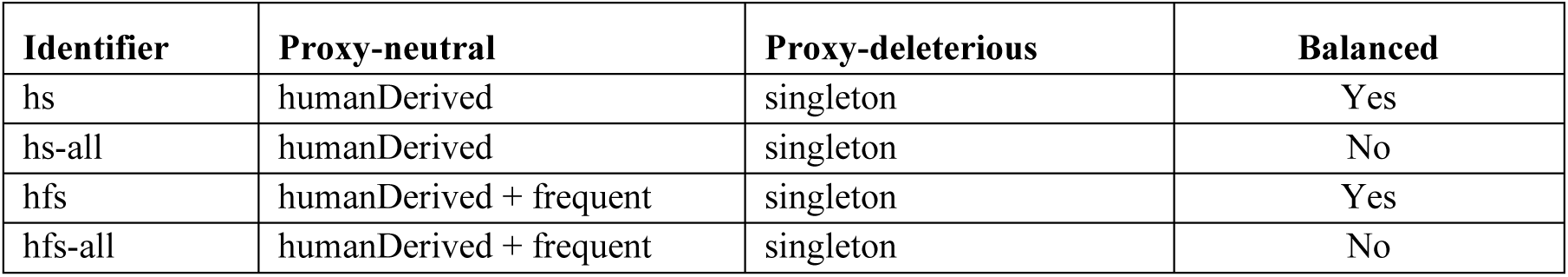
Overview of models with a combined set of training variants. For each combination, a balanced (with a sample of singleton variants) and an unbalanced (with the entire set of singleton variants) model was trained.

**Figure 4** shows the results on the ClinVar validation set (see Methods section *Model evaluation*). In terms of recall of pathogenic variants, the model hs (humanDerived-singleton) and hs-all (humanDerived-singleton-all) perform similarly (0.987 vs. 0.974), whereby the hs model has a slightly higher AUROC than CADD v1.6 (0.987 vs. 0.984). Contrary to our expectations, there is no improvement of the performance with the larger dataset containing all singleton variants. Instead, despite accounting for class imbalance during the training of the logistic regression model, the precision-recall score of the large dataset is notably lower than that of the balanced model and CADD v1.6. Conceivably, the large set of singletons introduces a significant amount of noise into the training data that outweighs the potential advantages of having larger training data.

**Figure 4.**
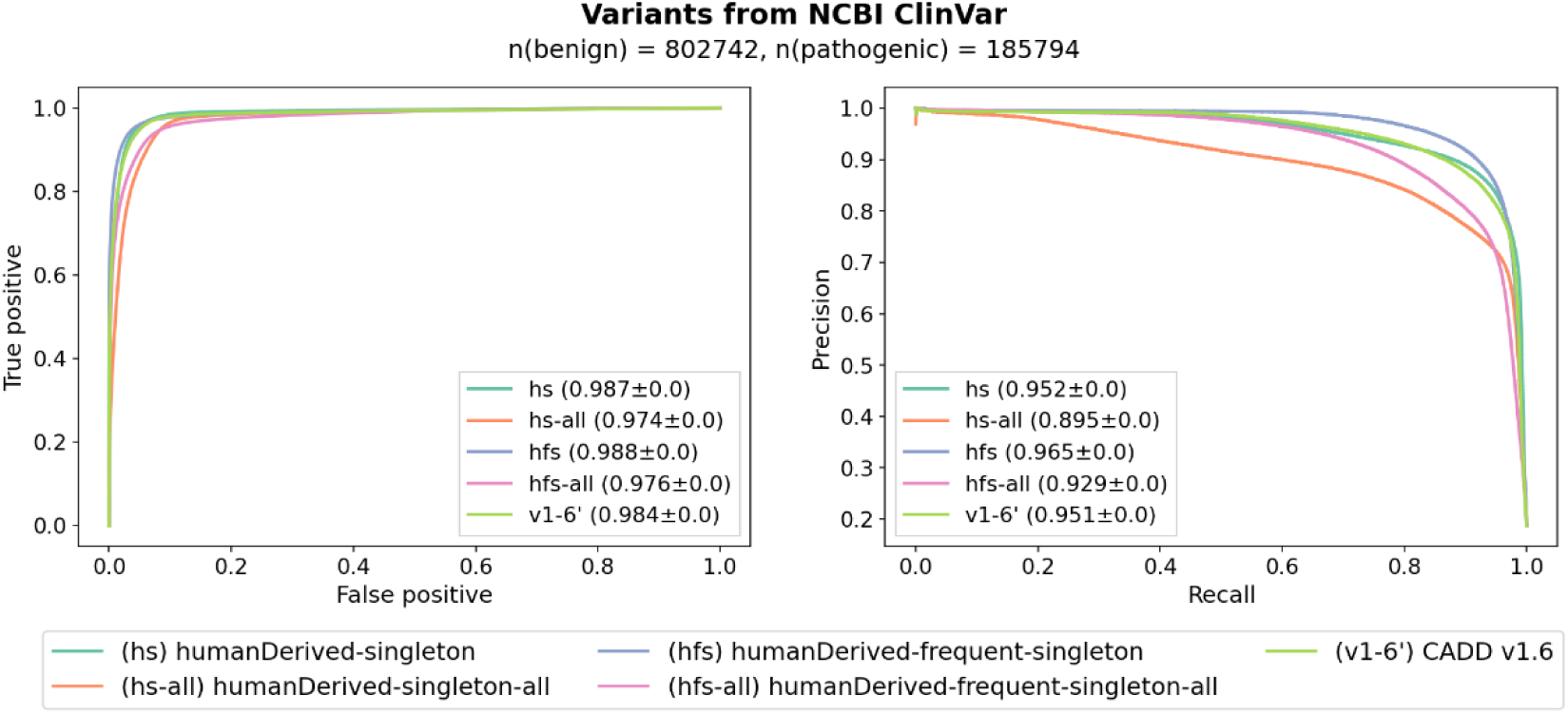
Performance of the alternative models trained on the combined sets of human-derived variants and standing variation. While the models trained on balanced datasets have comparable performance, the ones trained with the entire set of singleton variants perform worse. The model trained on the augmented dataset (hfs), i.e. on a combination of the human-derived variants and the frequent variation as a proxy-benign set and the singleton variants as a proxy-deleterious set, has the highest recall and precision of predictions compared to other alternative models as well as CADD v1.6.

### Augmenting human-derived variants with standing variation increases training data and improves performance

Large datasets of human variation may not only improve the original proxy-pathogenic set of CADD but may also allow for an extended proxy-benign set. The proxy-benign set of human-lineage derived fixed or high-frequency variants is based on variants that saw many generations of purifying selection, while being fixed among all humans or substantially increasing in frequency in humans. The purifying selection probably goes along with substantial shifts in functional constraint measures, like species conservation. Using these measures in distinguishing high and low effect variants might be advantageous in many cases, it may however miss effects of recent evolutionary changes that might also have an impact on fitness (Lupski et al. 2011). The analysis of model coefficients in the previous section showed that conservation and constraint features are less important when training models with standing variation, hence, including standing variation in the training set might be beneficial in terms of accounting for recent evolutionary changes. Thus, we train an additional model with an augmented set of proxy-benign variants, that contain a combination of human-derived and frequent variants and contrast it with a set of singleton variants.

Again, we train balanced (hfs for humanDerived-frequent-singleton) and unbalanced (hfs-all for humanDerived-frequent-singleton-all) models. For the balanced model, a subsample of the singleton set is used (40,267,173 SNVs and 9,033,4893 InDel). For the unbalanced one, all singleton variants (268,404,480 SNV and 27,934,609 InDel) are contrasted with the combined proxy-benign set. Note that all sequence differences in the original CADD proxy-benign training set were checked against the 1000 Genome database to exclude variants variable in the human population (requiring an allele frequency of more than 95%) (Kircher et al, 2014). As described before, models are trained for 50 iterations and compared using the ClinVar validation set.

**Figure 4** shows the performance of models compared with CADD v1.6. In terms of recall of pathogenic variants, we observe a slight improvement for the balanced model (hfs, 0.988) compared with CADD v1.6 (0.984) but deterioration for the unbalanced one (hfs-all, 0.976). Further, the precision of predictions of the balanced model has improved notably, having an AUPRC score of 0.965 compared to 0.951 of CADD v1.6. Apparently, inclusion of frequent standing variation improves the pathogenicity prediction of CADD scores. However, also here, a larger dataset of proxy-pathogenic variants (hfs-all) does not result in higher performance, presumably, due to introduction of noise from the very large set of singleton variants.

### Alternative models outperform in various variant groups

The ClinVar-based validation dataset used here seems saturated, i.e. the performance of most models is so high that selecting one best model is not possible. Thus, we assess the model performance by variant consequence, anticipating clearer differences between models for the more difficult to differentiate functional consequences levels. In this analysis, we include the previous models trained solely on standing variation, excluding those based on the rare-singleton set as they had the lowest performance across all consequence levels.

The evaluation of models delivers mixed results as there is no one model that outperforms all others across all or most variant groups (**Figure 5A**). Thus, CADD v1.6 is slightly better than alternative models in recall of nonsynonymous (0.787) and upstream variants (0.814), whereas, for example, the humanDerived-frequent-singleton model ranks first for inframe (0.850), 3’UTR (0.871) and 5’UTR (0.771) as well as stop-lost (0.806) consequences and the humanDerived-singleton model is better for synonymous (0.788), regulatory (0.776), splice-site (0.969) and non-coding change (0.942) variants. Most of the differences are small though and only range between 0.1 and a few percentage points.

**Figure 5.**
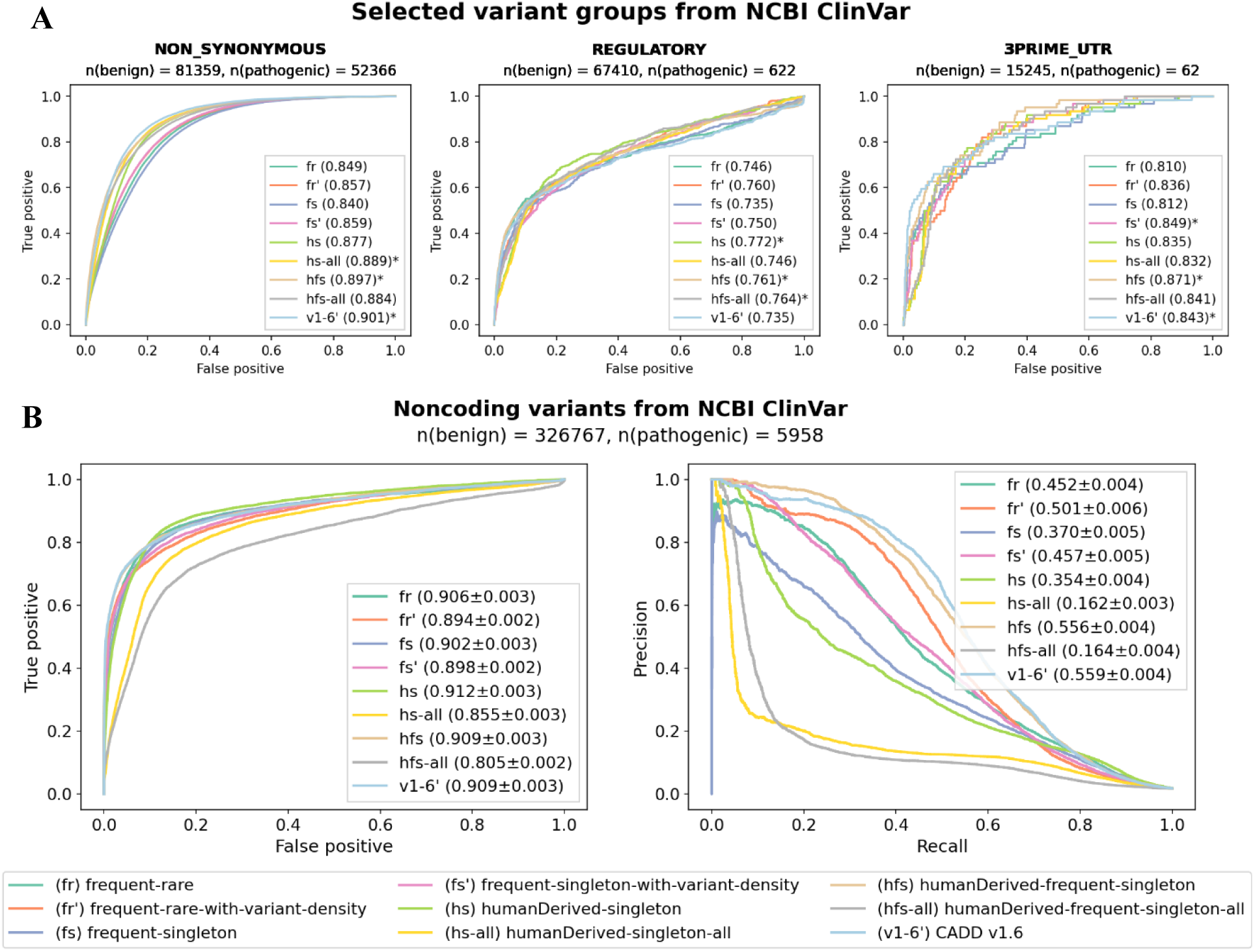
Model performance in various variant groups as well as on noncoding variants from NCBI ClinVar. A) Performance for selected groups of variant effects. Asterisks mark the models with the highest scores. Neither alternative models nor CADD v1.6 outperform other models in a majority of variant groups. Instead, each model has the highest recall rate in one or few groups. For example, CADD v1.6 has the highest recall for nonsynonymous, hs (humanDerived-singleton) for regulatory and hfs (humanDerived-frequent-singleton) for 3’UTRs variants. B) **Model performance aggregated over noncoding variants.** The differences of models become clearer when comparing them using a set of noncoding variants from the NCBI ClinVar database. While most of the models have comparable performance in terms of recall, a higher precision is only achieved by the hfs (humanDerived-frequent-singleton) and CADD v1.6 models.

In the next step, we analyze the validation set aggregated over coding (**Supplementary Figure 4**) and noncoding variants (**Figure 5B**) (for the definition of coding and noncoding regions see the Methods section on *Model evaluation*). The most notable differences between models can be observed for noncoding variants, whereas for coding positions, which make up most of the validation dataset with over 66% of all and 96% of the pathogenic variants, all models have rather similar performance. The large models containing the entire set of singleton variants have lower AUROC scores for predictions of noncoding variants (hs-all: 0.855, hfs-all: 0.805) and significantly lower AUPRC scores (hs-all: 0.162, hfs-all: 0.164). These models seem to assign higher deleteriousness scores to ClinVar variants, although their mean score on the 1 random million variants is in general lower than that of other models (**Error! Reference source not found.**). Interestingly, most of the alternative models have significantly lower precision-recall scores on noncoding variants compared to CADD v1.6. One exception is the model based on the combination of human-derived variants and standing variation (hfs). Not only has it comparable performance to CADD v1.6 on noncoding variants, but it also outperforms it in terms of prediction precision of coding variants (0.973 vs. 0.959). This suggests that the model benefits from augmentation of the CADD training dataset with frequent and singleton standing variation.

## Discussion

The prioritization of disease-causing variants with machine learning methods is of increasing importance for clinical genetics and precession medicine, making comprehensive and reliable genomic training data essential. Here, we presented several training sets based on human standing genetic variation and showed that they compete with a state-of-the-art model for genome-wide variant prioritization, CADD v1.6 that is based on evolutionary derived sequence changes and *de novo* variant simulations.

The idea to use human genetic variation was motivated by the observation that deleteriousness scores like CADD v1.6 are correlated with the observed allele frequency of variants in the population. This observation is in line with the rare disease model that argues that Mendelian diseases are caused by individually rare alleles of large effects. It is supported by evolutionary theory which predicts that disease causing variants are linked to reduced fitness and are selected against, so their frequency in the population should be low (Pritchard and Cox 2002; Bulmer 1971; Barton and Turelli 1989). At the same time, the presence of pathogenic variation in the human genome is explained by detrimental variants not being completely eliminated by purifying selection as mutations are being created in a large population, some diseases only causing modest effects on fitness, or late-onset diseases and recessive effects with only limited purifying selection (Gibson 2012). These considerations support the idea that leveraging variation in the human population can be beneficial for predicting deleteriousness.

While the application for genome-wide prediction of variant deleteriousness is novel, several methods have pioneered the integration of genetic variation data together with functional genomics for deriving measures of sequence constraint. For example, LINSIGHT integrates information on genetic polymorphisms within the human genome and its divergence from closely related species to assign a likelihood to noncoding genomic segments to have an impact on fitness (Huang, Gulko, and Siepel 2017). It relies on the idea that signals of past natural selection offer valuable insights into todays’ phenotypic importance. Another evolutionary based tool, PrimateAI, builds a deep neural network for pathogenicity assessment of missense variation using variants from gnomAD with an allele frequency of > 0.1% (Sundaram et al. 2018). The authors expanded a rather small set of common human missense variants with an extensive set of frequent missense variation observed in primates, showing that variants commonly observed in other primates are largely benign in humans. In this study, we build upon these previous observations to build genome-wide ML models of variant effects, showing that common human variation can be used as a proxy for benign allelic changes in all genomic regions, whereas (ultra-)rare variation can approximate deleterious variation in a binary classification approach. We show that this approach can compete with state-of-the-art methods for variant prioritization like CADD v1.6 and overcomes some limitations of conventional training data, like the artificial nature of simulated variants and a masking of human-derived changes in the calculation of sequence constraint measures.

Leveraging standing variation accounts for more recent evolutionary changes in the human genome, that are especially common in regulatory elements and that tend to be less conserved than, for example, protein-coding DNA (Danko et al. 2018; Suntsova and Buzdin 2020). Our results show that models based on standing variation put less focus on constraint and conservation measures during the model training and better utilize epigenetic and regulatory activity for the assessment of functional impact of variation. Standing variation offers a reasonable alternative for the simulated pathogenic data as it has a natural representation of variants across the functional elements compared to the simulated data, which only adjusts mutation rates in large genome windows. Further, datasets based on standing variation are already large, allowing a comprehensive sampling of feature values and in principle also allow for more complex models. The coverage of rare annotations and broad sampling of genetic regions is instrumental for better modeling of the underling biological processes in general and an improved variant prioritization in particular. The size of these datasets will continue to increase with an increasing number of sequenced genomes, improving the quality and accuracy of estimated allelic frequencies and enabling regression rather than classification methods.

The human-derived data used by CADD and other tools (Quang, Chen, and Xie 2015; Sundaram et al. 2018; Gao et al. 2023) offers a large dataset of likely benign genetic changes, that, however, needs elaborate curation of features and limits the choice of accessible annotations. For example, the conservation scores PhyloP and PhastCons must be recalculated for CADD, excluding the human reference sequence. This is necessary because human-derived variants would be represented in the human reference sequence and, consequently, these alignments and cause lower conservation scores when considered in calculation. With standing variation, conservation scores can be used off-the-shelf, i.e. as provided by UCSC Genome Browser and EBI ENSEMBL services. Another issue with human-derived data is that it is generated from the ancestral sequence’ perspective, i.e. the human reference allele is not the ancestral allele. Thus, several annotations must be switched in order to match the human reference perspective. For example, a stop-loss variant in the human-derived data must be annotated as stop-gain before model training. Consequently, the annotation of start loss and stop-gain variants is currently very limited. These and several other adaptations owing to the special origin of evolutionary data might lead to loss of information and introduce biases during training. Replacing human-derived data with standing variation overcomes these limitations, allowing to include new precalculated annotations into the training set.

Datasets based on standing variation also have some limitations. Rare and ultra-rare variants are used as an approximation for pathogenic variants; however, many rare variants will not have a detrimental effect on fitness. While the simulated pathogenic set of CADD also has an unknown rate of mislabeled variants (Rentzsch et al. 2019), its mislabeling is estimated to be 85%-95% - depending on the definition of functional constraint (Kellis et al. 2014). Estimation of the true proportion of deleterious rare variants is also challenging and varies significantly (Dukler et al. 2022). Another issue arises from potential sequence errors in the case of ultra-rare variants. In general, construction of databases of genomic variants is a complex process, where errors can occur and accumulate during each step including sequencing, mapping to the reference genome and variant calling (Nielsen et al. 2011; Li 2014; Pfeiffer et al. 2018). Extensive efforts have been made towards reduction of overall error rates like improving variant calling using recent advancements in machine learning (N.-C. Chen et al. 2023) or Telomere-to-Telomere sequencing of the genome (Nurk et al. 2022). Nonetheless, errors in databases of sequenced population data remain an important issue but might be compensated by the sheer number of rare variants and probably have limited impact on simple models.

A further drawback of the available variant data lies in the lack of ethnic diversity, which might negatively affect the allelic frequencies reported in datasets of genomic data. Out of 76,156 of sequenced genomes in gnomAD v3.0, almost 52% are of European and 27% of African ancestry (S. Chen et al. 2022). The new release of gnomAD v4.0 includes more than 400,000 exomes from the UK Biobank, where individuals of European ancestry account for over 94% of sequenced individuals (Nagar, Jordan, and Mariño-Ramírez 2023). It is known that, for example, African populations have higher genetic diversity and populations in general differ in the composition of common and rare variants (Witt et al. 2023; Choudhury et al. 2020). Thus, it is possible that rare or singleton variants in widespread databases of genomic data are common polymorphism of underrepresented populations. A solution here would be to access the allele frequencies on population level, assuming that a common benign variant in one population is benign in all populations.

The gnomAD v3.0 release used for this study excluded exome data, to obtain an even coverage of sequence variants along the genome and across functional effects. However, it might be argued that a call set including exome capture data increases the coverage of high impact functional effects and would improve model training, something that we did not explore here. We note though that CADD v1.6 outperforms most of the alternative models in coding regions, for example for synonymous and non-synonymous variant effects, where such a combination of exomes and genomes might give more weight to these molecular effects in training and thus improve the performance of models based on standing variation. The recently released gnomAD v4.0 release includes up to 730,947 exomes from various sources. Since the latest release available to us during the time of the study (v4.0) contained a known bug that affected allelic frequencies, we did not consider it for our study.

A general issue for benchmarking model performance of variant prioritization methods is the availability of validation sets. To compare model performance, we utilized the NCBI ClinVar database, which despite being broadly used, has several drawbacks. For one, like all available databases of pathogenic variants, it is biased towards the coding genome, thus containing an unproportionally small amount of noncoding variants, and even in the coding space it is biased towards well-studied genes (Landrum et al. 2018; Ellingford et al. 2022; Spielmann and Kircher 2022). So, if an alternative model has superior performance in noncoding genomic regions, the power to detect it on this validation set is limited. Secondly, the benign variants contained in the ClinVar dataset can be regarded only as conditionally benign as many of them were initially candidate disease variants or were at least prioritized with a clinical analysis pipeline, and later labeled as benign or Variant of Uncertain Significance (VUS) after obtaining no confirmation of their deleteriousness. This, nevertheless, does not mean that these ‘benign variants’ cannot be causative for any other genetic disease, i.e. ‘absence of evidence’ does not necessarily mean ‘evidence of absence’ (Ciesielski et al. 2024). To overcome this issue, many studies including CADD v1.6 use common variation with MAF > 1% or 5% from population studies as a benign set and contrast them with the pathogenic variation from ClinVar to validate results. However, since common variation is a part of some of the alternative training sets, we did not follow this approach to avoid inflated performance of models based on frequent variants.

## Conclusions

Here, we present several training sets based on standing variation from gnomAD v3.0, approximating variant deleteriousness with low or high allele frequency in the population. We show that using frequent standing variation as a proxy for benign and rare and singleton variation as a proxy for deleterious variants allows to train state-of-the-art models for genome-wide variant prioritization. The proposed datasets have several advantages, like being significantly larger and potentially less biased than conventional datasets for deleteriousness scoring. Consequently, some of the alternative models outperform a broadly used score for variant prioritization, CADD v1.6, in predictions of pathogenic and benign variants from the NCBI ClinVar database.

Covering a broader range of rare annotations and containing recent evolutionary changes, large datasets of standing variation can be used either standalone or in combination with evolutionary data, for example human-derived variants or primate variation data, for various research purposes. Across various training data settings, we show that these alternative datasets can improve variant prioritization, providing better assessment of variant pathogenicity especially also for non-coding effects. We make these training sets readily available for reproducing our results on the CADD v1.6 feature set, but also for annotating them with other feature sets. Future models derived from the variants sets provided will enable various basic research and clinical applications.

## Methods

### Correlation of allele frequencies and deleteriousness scores

CADD v1.6 scores for gnomAD v3.0 SNVs were downloaded from the CADD webserver (https://kircherlab.bihealth.org/download/CADD/v1.6/GRCh38/gnomad.genomes.r3.0.snv.tsv.gz). Their allele frequency (AF) was looked up in the gnomAD v3.0 file downloaded from the UCSC Genome Browser server (https://hgdownload.soe.ucsc.edu/gbdb/hg38/gnomAD/vcf/gnomad.genomes.r3.0.sites.vcf.gz) using tabix (Li 2011). For each of the fifty bins of PHRED-scaled CADD scores in the interval (0, 50], 100,000 variants were selected, excluding variants with zero allele frequency (AF=0). For PHRED scores up to 29, it was possible to extract the intended number of positions, but for variants with a PHRED score 30 and higher, the number of available positions is decreasing, so that, for example, for the bin (49,50] only 581 variants were available. In total, 3,264,650 positions were retrieved. Data was plotted with mean allele frequency values for each of the fifty bins using Python package *seaborn v0.13.2*. Additionally, the number of singletons in each bin was plotted where singletons are defined as variants with allele frequency equaling one (AF=1) in the gnomAD v3.0 file. Spearman correlations of allele frequencies and PHRED-scaled CADD scores were measured across concatenated value pairs for all bins (n=3,264,650) using *scipy v1.9.0* package.

### CADD training dataset

CADD training data is composed of equally large proxy-pathogenic and proxy-benign sets of SNVs and InDels. The proxy-benign set contains evolutionary derived sequence alterations (fixed or at very high frequency on the human-lineage) that saw many generations of purifying selections (Kircher et al. 2014). The proxy-pathogenic set is produced by a “germline simulator” of *de novo* variants that accommodates some properties of the proxy-benign variants such as substitution rates and mutations in a CpG context. Detailed information can be found in the original CADD publications (Kircher et al. 2014). The feature set of CADD v1.6 (Rentzsch et al. 2021) contains over 100 different annotations of sequence conservation and constraint, epigenetic and regulatory activity, variant density and others. The full list of annotations can be found in the respective CADD v1.6 release notes (https://cadd.bihealth.org/static/ReleaseNotes_CADD_v1.6.pdf).

### gnomAD variants and dataset matching

gnomAD v3.0 variants were downloaded from the UCSC Genome Browser (https://hgdownload.soe.ucsc.edu/gbdb/hg38/gnomAD/vcf/gnomad.genomes.r3.0.sites.vcf.gz) and SNVs and InDels were selected according to the allele frequencies thresholds defined in Table 1. Variants with an average AF > 0.5 across all individuals were not considered. For training of models based solely on standing variation, the rare and singleton variants were subsampled to match the number of variants in the frequent set, in order to have balanced training data with equal number of proxy-benign and proxy-pathogenic variants. Thereby, the substitution rates as well the length of insertions and deletions were matched to that in the frequent set to avoid label leakage.

For large models containing the entire set of singletons (hs-all, hfs-all), training datasets were not matched in their composition of substitution rates or length of insertions and deletions to keep the largest possible number of variants. The dataset for the hs model was sampled from the matched singleton set so that it is matched to the frequent set. Singletons for the hfs model were matched to the combined set of human-derived and frequent variants. The analysis of resulting model coefficients reveals that features containing the reference and alternative alleles or their length are of little importance for model predictions. Consequently, the exact matching seems of limited importance. All variant sets are annotated with the full set of annotations derived from CADD v1.6, including variant density.

### Model training

All alternative models are trained using the CADD framework. In the first step, the variant sets are annotated all annotations used for CADD v1.6 training. In the next step, the framework imputes the annotated variants by filling missing values, binarizes categorical features and creates feature crosses (multiplication of selected features) of select annotations. In the imputation step, annotations of variant density are omitted for models trained without those features. This results in 1028 features for the full set of annotations and 848 features without variant density annotations. The features in each dataset are normalized and used for the training of a logistic regression model using version 0.20.4 of *scikit-learn*. In the first part of the analysis, all models are trained for 20 iterations. In the second part of the analysis the models that consist of a combination of human-derived variants and standing variation are trained for 50 iterations. Regularization parameters remained unchanged from CADD v1.6.

### Validation sets

The validation sets used to identify the optimal iteration of logistic models are described in **Error! Reference source not found.**. These are datasets previously described for CADD model evaluation (Rentzsch et al. 2019; Schubach et al. 2024) and were chosen to cover a wide range of variant effects from pathogenicity to molecular impact. Since the datasets vary significantly in size, an unweighted average was used for model tuning (i.e. the choice of the optimal number of iterations).

The validation set used to compare trained models with CADD v1.6 is based on the NCBI ClinVar database, as accessed in June 2023 (https://ftp.ncbi.nlm.nih.gov/pub/clinvar/tab_delimited/archive/variant_summary_2023-06.txt.gz). The dataset was filtered for single nucleotide variants as well as insertion and deletions of no longer than 50 base pairs mapped to the GRCh38 genome build. Variants with clinical significance ‘Pathogenic’, ‘Likely pathogenic’, ‘Pathogenic/Likely pathogenic’, ‘Likely benign’, ‘Benign’ and ‘Benign/Likely benign’ were selected. Variants located on the mitochondrial genome were dropped from the analysis. The resulting dataset of 988,536 variants (185,794 pathogenic and 802,742 benign) was annotated using CADD v1.6 adding a column ‘Consequence’ based on the Ensembl VEP annotation build v95 (McLaren et al. 2016). This column was used for model comparisons on consequence level as well as for the separation of data into coding and noncoding sets (coding being defined as ‘STOP_GAINED’, ‘STOP_LOST’, ‘CANONICAL_SPLICE’, ‘NON_SYNONYMOUS’, ‘SYNONYMOUS’, ‘FRAME_SHIFT’, ‘INFRAME’ and noncoding being defined as ‘INTRONIC’, ‘REGULATORY’, ‘SPLICE_SITE’, ‘3PRIME_UTR’, ‘NONCODING_CHANGE’, ‘DOWNSTREAM’, ‘UPSTREAM’, ‘5PRIME_UTR’, ‘INTERGENIC’).

### Model evaluation

The models are evaluated using the Python *scikit-learn v1.1.1* package (metrics: RocCurveDisplay, PrecisionRecallDisplay, roc_curve, auc, average_precision_score) and visualized with *seaborn v0.13.2* and *matplotlib v3.7.0*. Pearson and Spearman correlations as well as significance levels were calculated with *scipy v1.9.0*.

### Visualization of model coefficients

As the input data for the training of the logistic model is normalized (mean 0, standard deviation of 1), coefficients can be directly interpreted as feature importance. The 30 most important features (with the highest absolute value of the coefficient) were selected for each model and visualized in a stacked bar plot. The size of each block in a bar represents the absolute value of a model coefficient, the total length of a bar indicates the sum of coefficients. To facilitate the comparison, the features were grouped into five categories according to **Error! Reference source not found.**.

## Abbreviations

CADD: Combined Annotation Dependent Depletion
AUROC: Area Under Receiver Operating Characteristic
AUPRC: Area Under Precision Recall Curve
gnomAD: Genome Aggregation Database
MAF: Minor Allele Frequency
SNV: Single Nucleotide Variant
InDel: insertion/deletion

## Declarations

### Ethics approval and consent to participate

Not applicable

### Consent for publication

Not applicable

### Availability of data and materials

Annotated training data sets as well as model coefficients are available on Zenodo.

### Authors’ contributions

P.R. created the datasets of standing variation and trained models from the first part of the manuscript.

L.N. created the datasets and trained the models for the second part, analyzed the results of the first and second parts, wrote the manuscript with the support of M.K. and P.R. M.K and P.R. supervised the project. All authors discussed the results and commented on the manuscript.

## Acknowledgements

We thank current and previous members of the CADD project and Kircher lab for helpful discussions and suggestions. We especially thank GM Cooper and J Shendure for their ideas and contributions to the development of CADD. Computation has been performed on the HPC for Research cluster of the Berlin Institute of Health at Charité – Universitätsmedizin Berlin as well as on the OMICS cluster at the University of Lübeck.

