## Supplementary Materials for "varCADD: large sets of standing genetic variation enable genome-wide pathogenicity prediction"

#### Supplementary Figures

##### Supplementary Figure 1

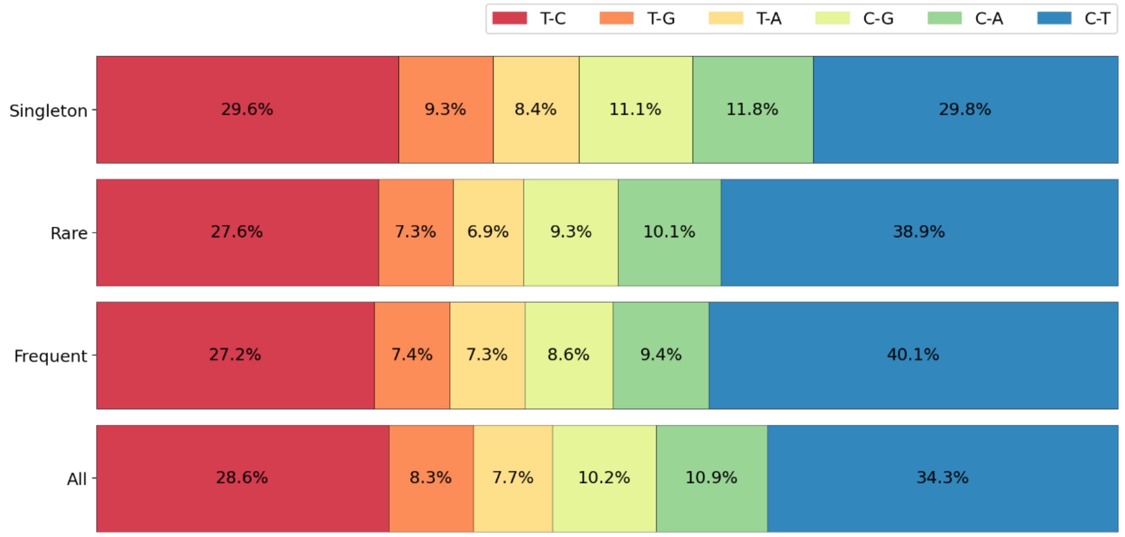


***Supplementary Figure 1: Selected SNV substitution frequencies in different training set groups.*** *Frequencies are based on a sample of ten million single nucleotide variants. The substitution rates differ for different groups of allele frequencies and derived training set definitions. Thus, for example, the C to T substitution accounts for 40.1% in the frequent group and only for 29.8% in the singleton group. As the reference and alternative alleles as well as transition/transversion state are features in the training set, the substitution frequencies have been matched to avoid label leakage during training.*

#### Supplementary Figure 2

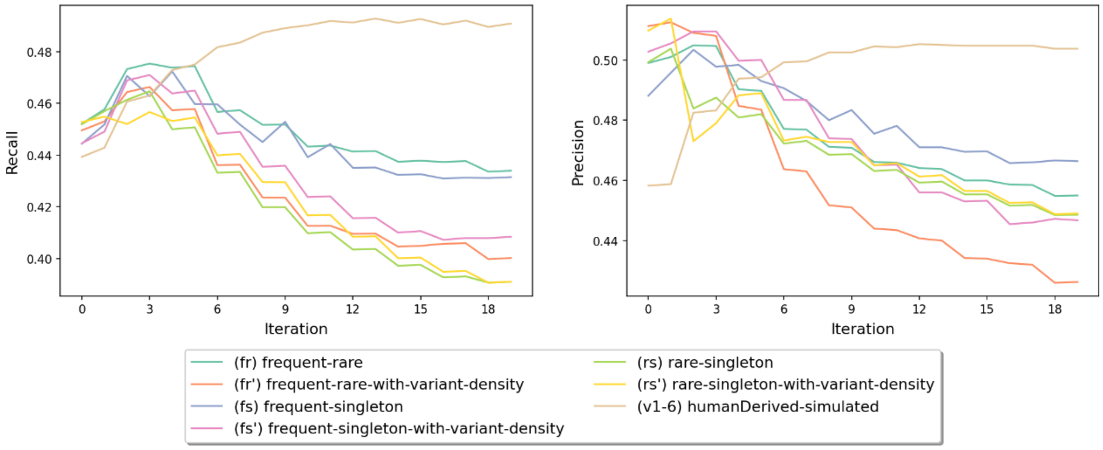


***Supplementary Figure 2: AUROC and AUPRC values of trained models depending on training iterations.*** *Models are evaluated on 10 validation sets (Supplementary Table 1) and AUROC and AUPRC values are averaged. We used unweighted averages to avoid larger datasets dominating the results, as the number of variants in each validation set varies significantly. The best number of iterations selected for the alternative models differs from that of CADD v1.6 (13$^{th}$ iteration). All models based on standing variation achieve their optimal performance rather early, after which performance drops significantly. This might be an indication of overtraining with increased number of iterations, lowering generalization power of the models when tested on validation sets. Interestingly, this holds true for all models, i.e. with and without annotations of variant density. Further, the only trained model that shows increasing AUROC and AUPRC is the one based on the original CADD training set without annotations of variant density. It however converges around the 10$^{th}$ iteration.*

#### Supplementary Figure 3

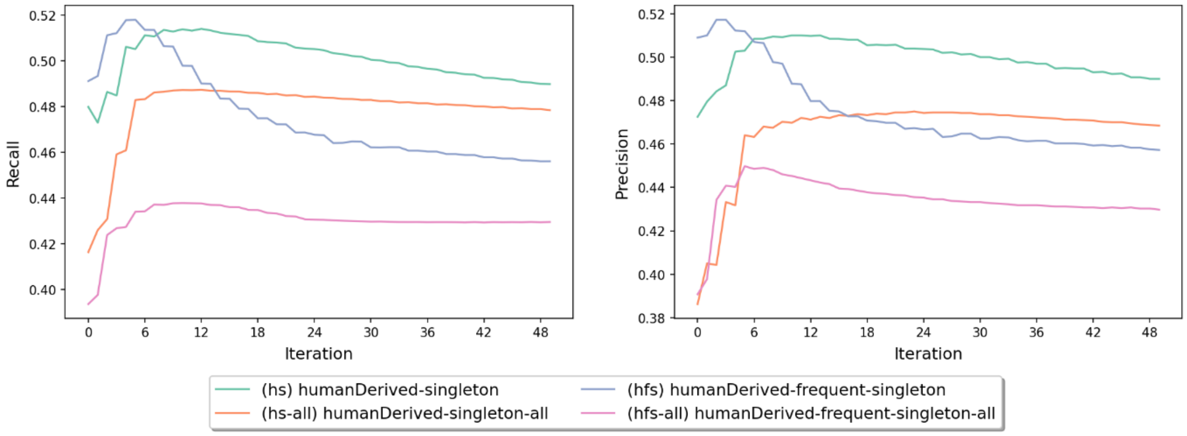


***Supplementary Figure 3: AUROC and AUPRC values of models based on different training iterations for models trained on the combination of human-derived data and standing variation.*** *To choose the optimal iteration non-aggregated performance measurements were also considered, but here the unweighted average across all validation sets is shown as described before. All models achieve the optimal performance rather early in training, after which it decreases, i.e. models do not achieve a stable performance for more training iterations, but potentially overtrain on the surrogate training task.*

#### Supplementary Figure 4

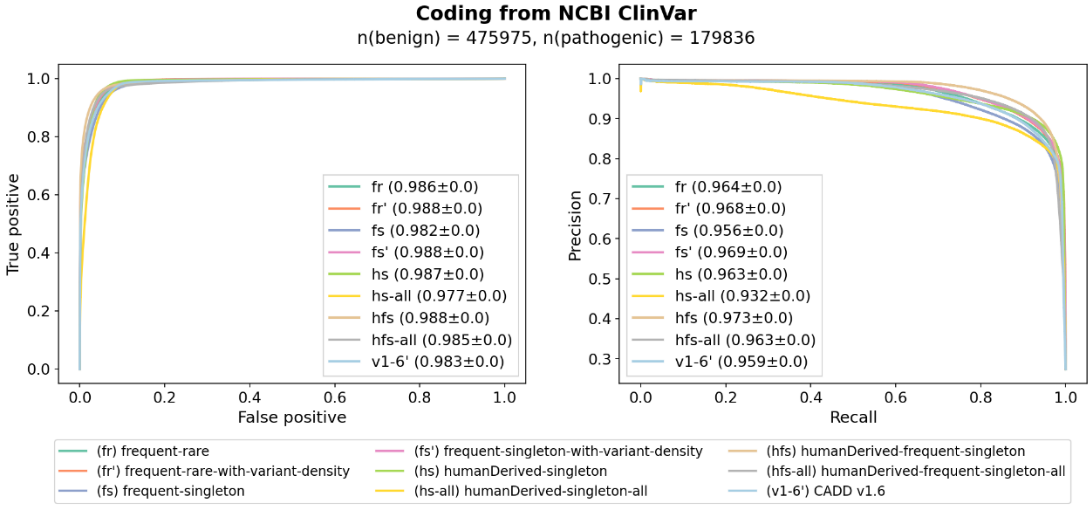


***Supplementary Figure 4. AUROC and AUPRC of models for coding variants from NCBI ClinVar.*** *All models perform similarly in terms of recall of coding variants, slightly bigger differences can be observed for the precision of model predictions. Unbalanced models trained with the entire set of singleton variants tend to have lower AUPRC scores, which indicates that class imbalance worsens model performance despite being considered as different class-weights during training. The hfs (humanDerived-frequent-singleton) model has the best performance when compared with other alternative models or with CADD v1.6.*

#### Supplementary Figure 5

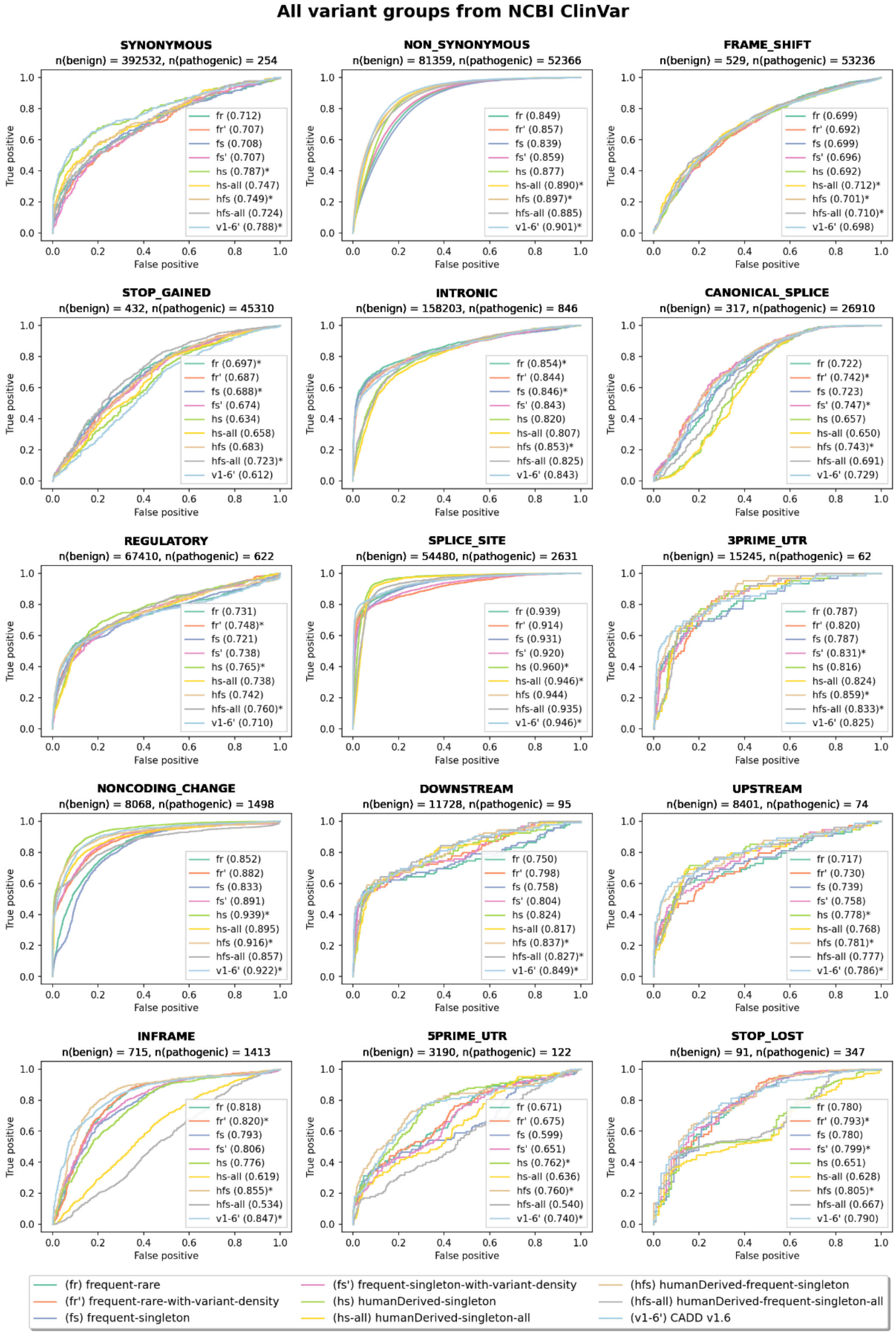


***Supplementary Figure 5: AUROC of models across variant groups.*** *The best models in each group are indicated with an asterisk. In general, no one model outperforms all others in recalling pathogenic variants from the NCBI ClinVar dataset but in each group, the best performing models are different. Consequently, CADD has high recall rates for synonymous, non-synonymous and stop-lost variants, whereas for example the hfs (humanDerived-frequent-singleton) model ranks first in recalling pathogenic 3' and 5' UTR variants.*

#### Supplementary Figure 6

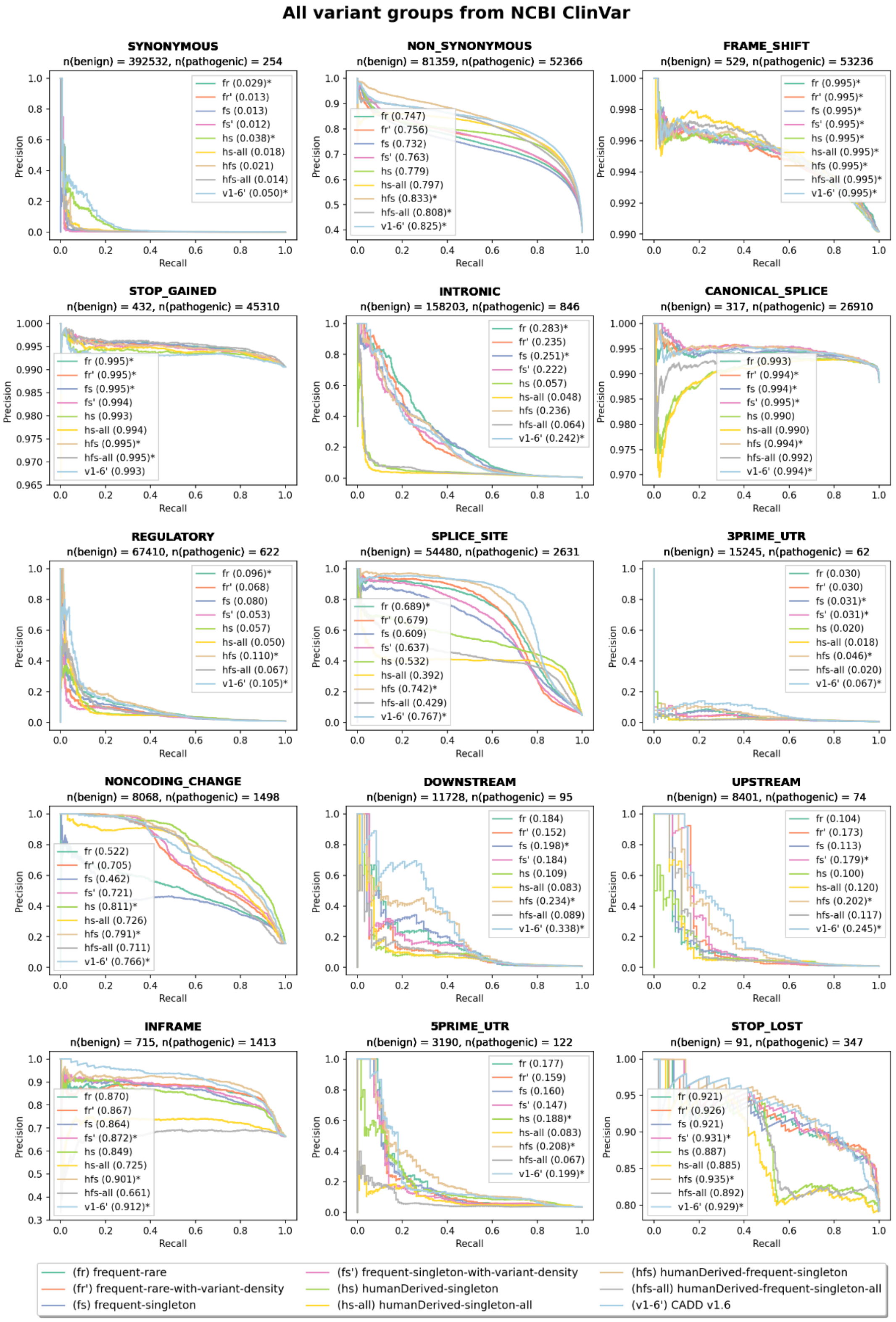


***Supplementary Figure 7: AUPRC of models across variant groups.*** *The best models in each group are indicated with an asterisk. Precision of models by variant consequence varies significantly. For example, frame-shift variants are identified by all models with a precision-recall score of 0.995, whereas the highest precision on synonymous variants is 0.059, which indicates that synonymous variants are hard to classify as benign or pathogenic. On the contrary to recall scores, two models, CADD v1.6 and hfs (humanDerived-frequent-singleton), outperform others in terms of precision of predictions.*

#### Supplementary Figure 7

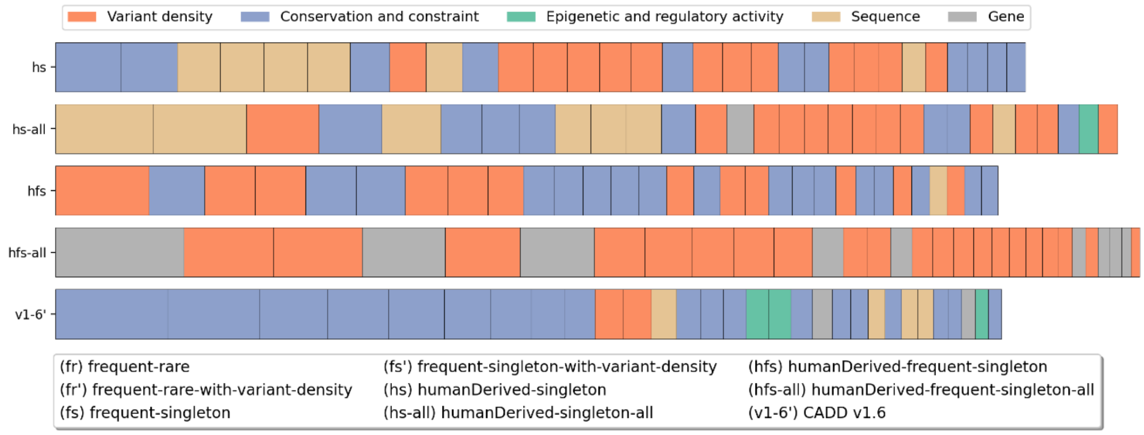


***Supplementary Figure 8: Model coefficients of the first 30 most important features or feature combinations of models based on a combination of human-derived data and standing variation.*** *The size of each block represents the absolute value of the model coefficients, the total length of a bar indicates the sum of coefficients. As all models contain annotations of variant density, they play a significant role for models trained with standing variation but to a lesser extent than for models based solely on standing variation (**Error! Reference source not found.D**). Instead, species conservation and other constraint measures are more important here, which may be attributed to the presence of human-derived variants in the training set.*

### Supplementary Tables

#### Supplementary Table 1

| Name | Size | Proportion of Benign (0) and pathogenic (1) | Source |
|---|---|---|---|
| FANC | 2770 | 0: 0.88, 1: 0.12 | (Rogers et al. 2014) |
| MFASS-exon | 14130 | 0: 0.97, 1: 0.03 | (Chong et al. 2019) |
| MFASS-intron | 13603 | 0: 0.96, 1: 0.04 | (Chong et al. 2019) |
| ClinVar-full-ExAc | 105643 | 0: 0.48, 1: 0.52 | Pathogenic: (Landrum et al. 2018)<br>Benign: (Lek et al. 2016) |
| BRCA1-E3 | 1355 | - | (Findlay et al. 2018) |
| YAP1-single | 180 | - | (Gray et al. 2018) |
| DLG4-single | 500 | - | (Gray et al. 2018) |
| sM-TERT-GBM | 777 | - | (Kircher et al. 2019) |
| sM-LDLR | 954 | - | (Kircher et al. 2019) |
| sM-HBB | 210 | - | (Patwardhan et al. 2009) |

***Supplementary Table 1: Validation sets used to tune the number of training iterations of the alternative models.*** *The validation sets include manually curated data such as ClinVar pathogenic versus common human genetic variants or experimental readouts from saturation mutagenesis experiments of non-coding sequence elements.*

#### Supplementary Table 2

| Model abbreviation | Model | Optimal iteration |
|---|---|---|
| fr | frequent-rare | 4 |
| fr’ | frequent-rare-with-variant-density | 3 |
| fs | frequent-singleton | 3 |
| fs’ | frequent-singleton-with-variant-density | 3 |
| rs | rare-singleton | 4 |
| rs’ | rare-singleton-with-variant-density | 4 |
| hs | humanDerived-singleton | 8 |
| hs-all | humanDerived-singleton-all | 6 |
| hfs | humanDerived-frequent-singleton | 5 |
| hfs-all | humanDerived-frequent-singleton-all | 6 |
| v1-6 | humanDerived-simulation | 11 |
| v1-6’ | CADD v1.6 | 13 |

***Supplementary Table 2: Optimal number of training iterations of all models based on the validation sets.*** *Most models trained with data on standing variation achieve the optimal performance early, including those trained on the entire set of singleton variants (hs-all and hfs-all). The original CADD v1.6 achieves its best performance around the $13^{th}$ iteration, which is the value previously used for this version. A retrained version without the variant density features (v1-6) performs best for only 11 iterations.*

#### Supplementary Table 3

| Group | All following features as well as combination with those |
|---|---|
| Variant density | Sngl\|rare\|freq |
| Conservation and constraint | phylop\|gerp\|phcons\|bstatistic |
| Epigenetic and regulatory activity | encode\|chmm_\|ensembleregulatoryfeature |
| Sequence | type\|ref\|length\|toverlapmotifs\|motifdist\|alt\|remapoverlaptf\|remapoverlapcl\|nuccomb\|cpg\|gc\|remapoverlaptf |
| Gene | sift\|spliceai\|polyphen |

***Supplementary Table 3: The groups of features used for analysis of model coefficients.*** *The feature groups were created using feature names that contain the according strings. For example, the group variant density contains features like priphcons, mamphcons, verphcons, priphylop, mamphylop, verphylop, gerpn, gerps, gerprs, gerprspval, bstatistic as well as features crosses that contain one of those, e.g. (3PRIME_UTR, bstatistic) or (NON_SYNONYMOUS, mamphylop). The table is used to identify only the 30 most important features; therefore, it might miss a proportion of relevant feature contributions to each full model.*

#### Supplementary Table 4

| | fr | fr' | fs | fs' | rs | rs' | hs | hs-all | hfs | hfs-all | v1-6 | CADD v1.6 | Mean | Variance |
|---|---|---|---|---|---|---|---|---|---|---|---|---|---|---|
| **fr** | 1.0 | 0.713 | 0.978 | 0.786 | 0.925 | 0.831 | 0.674 | 0.392 | 0.74 | 0.401 | 0.687 | 0.65 | 0.507 | 0.037 |
| **fr'** | 0.713 | 1.0 | 0.695 | 0.979 | 0.647 | 0.631 | 0.565 | 0.36 | 0.869 | 0.678 | 0.496 | 0.492 | 0.568 | 0.061 |
| **fs** | 0.978 | 0.695 | 1.0 | 0.794 | 0.972 | 0.908 | 0.642 | 0.349 | 0.737 | 0.378 | 0.674 | 0.633 | 0.534 | 0.059 |
| **fs'** | 0.786 | 0.979 | 0.794 | 1.0 | 0.765 | 0.768 | 0.604 | 0.372 | 0.905 | 0.656 | 0.571 | 0.555 | 0.571 | 0.061 |
| **rs** | 0.925 | 0.647 | 0.972 | 0.765 | 1.0 | 0.958 | 0.608 | 0.328 | 0.706 | 0.359 | 0.648 | 0.606 | 0.501 | 0.023 |
| **rs'** | 0.831 | 0.631 | 0.908 | 0.768 | 0.958 | 1.0 | 0.568 | 0.305 | 0.724 | 0.382 | 0.623 | 0.579 | 0.501 | 0.019 |
| **hs** | 0.674 | 0.565 | 0.642 | 0.604 | 0.608 | 0.568 | 1.0 | 0.867 | 0.751 | 0.703 | 0.864 | 0.906 | 0.444 | 0.091 |
| **hs-all** | 0.392 | 0.36 | 0.349 | 0.372 | 0.328 | 0.305 | 0.867 | 1.0 | 0.62 | 0.84 | 0.737 | 0.815 | 0.477 | 0.097 |
| **hfs** | 0.74 | 0.869 | 0.737 | 0.905 | 0.706 | 0.724 | 0.751 | 0.62 | 1.0 | 0.785 | 0.793 | 0.779 | 0.562 | 0.082 |
| **hfs-all** | 0.401 | 0.678 | 0.378 | 0.656 | 0.359 | 0.382 | 0.703 | 0.84 | 0.785 | 1.0 | 0.585 | 0.653 | 0.488 | 0.077 |
| **v1-6** | 0.687 | 0.496 | 0.674 | 0.571 | 0.648 | 0.623 | 0.864 | 0.737 | 0.793 | 0.585 | 1.0 | 0.973 | 0.558 | 0.105 |
| **CADD v1.6** | 0.65 | 0.492 | 0.633 | 0.555 | 0.606 | 0.579 | 0.906 | 0.815 | 0.779 | 0.653 | 0.973 | 1.0 | 0.561 | 0.107 |

***Supplementary Table 4: Correlation of model scores on one million randomly sampled variants as well as the mean and the variance of scores.*** *The model trained on the humanDerived-singleton set of variants has the highest correlation with CADD v1.6 scores (except for a model trained on the original CADD data but without annotations of variants density, called v1-6 here). This is an indication that the simulated pathogenic set and the singleton set are similar but still have some differences so that models produce different scores. Interestingly, models with overall good performance like CADD v1.6, hfs, fr' and fs' have relatively high score averages, whereas models hs and hs-all have lower mean values (0.444 und 0.477).*
